# An Investigation of the YidC-Mediated Membrane Insertion of Pf3 Coat Protein Using Molecular Dynamics Simulations

**DOI:** 10.1101/2022.05.28.493840

**Authors:** Adithya Polasa, Jeevapani Hettige, Kalyan Immadisetty, Mahmoud Moradi

**Author notes:** Pacific Northwest National Laboratory, Richland, WA 99352. St. Jude Children’s Research Hospital, Memphis, TN 38105.

## Abstract

YidC is a membrane protein that facilitates the insertion of newly synthesized proteins into lipid membranes. Through YidC, proteins are inserted into the lipid bilayer via the SecYEG-dependent complex. Additionally, YidC functions as a chaperone in protein folding processes. Several studies have provided evidence of its independent insertion mechanism. However, the mechanistic details of the YidC independent protein insertion mechanism remain elusive at the molecular level. This study elucidates the insertion mechanism of YidC at an atomic level through a combination of equilibrium and non-equilibrium molecular dynamics (MD) simulations. Different docking models of YidC-Pf3 in the lipid bilayer were built in this study to better understand the insertion mechanism. To conduct a complete investigation of the conformational difference between the two docking models developed, we used classical molecular dynamics simulations supplemented with a non-equilibrium technique. Our findings indicate that the YidC transmembrane (TM) groove is essential for this high-affinity interaction and that the hydrophilic nature of the YidC groove plays an important role in protein transport across the cytoplasmic membrane bilayer to the periplasmic side. At different stages of the insertion process, conformational changes in YidC’s TM domain and membrane core have a mechanistic effect on the Pf3 coat. Furthermore, during the insertion phase, the hydration and dehydration of the YidC’s hydrophilic groove are critical. These demonstrate that Pf3 interactions with the membrane and YidC vary in different conformational states during the insertion process. Finally, this extensive study directly confirms that YidC functions as an independent insertase.

## Introduction

Approximately 33% of all membrane proteins are inserted and embedded in the plasma membrane bilayer during co-translation.^1,2^ The membrane proteins YidC/Oxa1/Alb3 work to fold incoming peptides into the membrane as efficiently as possible.^3–7^ YidC catalyzes the transmembrane insertion of newly synthesized membrane proteins in the absence of an energy supply domain, such as an ATPase,^8^ and is also involved in the insertion and placement of membrane proteins in microbes. The extent to which insertase proteins are required for inserting proteins into the membrane has been thoroughly investigated. They can be found in all aspects of life and are necessary for cell viability.^9–12^ They are adaptable proteins and can function along with the SecYEG pathway to insert peptides into the membrane through the Signal Recognition Particle (SRP) mechanism. They can fold and insert proteins independently of the Sec pathway.^9,13–19^ This study primarily focuses on the conformational dynamics of YidC, including both local and global conformational changes involved in the insertion process of the Pf3 coat protein.

YidC completes its function either independently as a membrane insertase or as a chaperone in the SecYEG complex mechanism. Additionally, YidC plays a critical role in the insertion of the LacY lactose permease membrane layer protein. ^15,20–22^ Also, YidC is involved in the incorporation of subunit II of cytochrome o oxidase in *E*.*Coli*.^23–25^ Initially, the Sec-autonomous pathway was thought to function without the contribution of an insertase. However, many studies have demonstrated that YidC is fundamental for the addition of small phage coat proteins like Pf3 coat and M13 in a Sec-free pathway. ^9,26–31^

A few experimental studies have explored the role of YidC in various microbial creatures. The genomes of most gram-positive microscopic organisms encode two YidC proteins: YidC1 and YidC2.^32,33^ Although YidC typically exists as a dimer or tetramer^34^ under physiological conditions, it was discovered that YidC can also function as a monomer in lipid bilayers.^8^ The protein is firmly anchored in the lipid bilayer by interfacial aromatic residues, a cytoplasmic salt-bridge group, and a periplasmic helix enhanced with aromatic residues. A group of aromatic residues above R72 may provide a polar hydrophobic surface with managing peptide insertion into the lipid bilayer. ^35,36^ The C–terminus of monomeric YidC cooperates with the ribosomes, and the short interhelical loops come into contact with the ribosomal proteins.^37^ YidC is believed to promote membrane insertion simply by binding nascent chains and promoting their insertion into the lipid bilayer using hydrophobic force. ^8^ The hydrophilic groove inside the membrane core of the YidC will increase the rate of accepting the hydrophilic moieties of a substrate into the membrane. ^38,39^ The YidC hydrophilic region traverses the inner side of the membrane and is closed to the periplasmic side of the bilayer. This decreases the hydrophobicity of the membrane towards the external side of the lipid bilayer. This hypothesis of the YidC mechanism provides excellent knowledge to study the conformational dynamics of YidC. In the first step, it interacts with a hydrophilic protein region temporarily in its groove, and in the second step, this peptide is translocated to the periplasmic side.^17^

Many prior studies have reported various explanations of the YidC independent mechanism. However, the global and local structural changes that occur in YidC during the process are not completely defined. It’s unknown how the cytoplasmic hairpin region of YidC and the water molecule inside the groove area of YidC take action during the insertion process. How would the incoming peptide or protein’s structure and conformation change during the process? We examined this topic using a combination of docking, classical molecular dynamics, and non-equilibrium simulations to analyze Sec-independent YidC’s (Fig. 1) insertion of the Pf3 coat transitionally into the membrane. In this study, we looked at the local and global conformational changes of YidC associated with Pf3 insertion into the hydrophilic groove, Pf3 interactions with YidC and the membrane, and conformational changes in Pf3 that occurred during the insertion process.

**Fig. 1.**
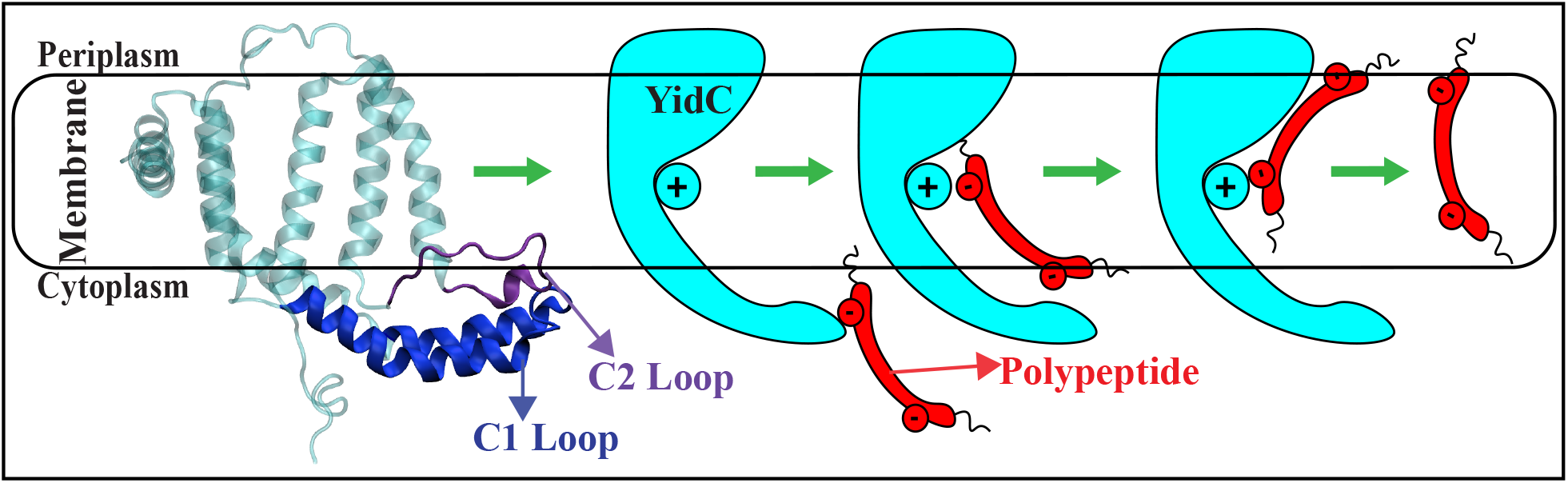
The cartoon representation of YidC and the schematic illustration of the secindependent insertion mechanism. A cartoon representation of YidC’s C1 and C2 loops on the cytoplasmic side (left). Schematic illustration of the YidC Sec-independent insertion of polypeptide (right).

## Methods

The crystal structure of YidC (PDB:3WO7^40^) was downloaded from the Protein Data Bank. Initially, the system was prepared using the Molecular Operating Environment (MOE) software^41^ by removing the water molecules from the crystal structure and assigning the appropriate protonation states for the residues using the protonate3D facility. We used MOE software to create two docking structures of the Pf3 coat protein interacting with YidC based on the previously hypothesized stages involved in the YidC sec-independent insertion process.^8,42^ In pose1, Pf3 is docked in the YidC’s hydrophilic groove (Fig. 2A) to evaluate probable interactions and conformational changes in the mechanism’s initial phase. The Pf3 coat is docked near to the periplasmic side (Fig. 2A) of the protein in pose2 to identify the interactions and conformational changes involved in the mechanism’s final phase. Biased and unbiased all-atom molecular dynamics (MD) simulations were performed to characterize the conformational differences of the two bacterial YidC2 - Pf3 docking models pose1 and pose2 (Fig. 2A) in a membrane environment. All simulations were performed with the NAMD 2.13^43^ using the CHARMM36m^44^ force field.^45^ TIP3P^46^ waters were used to solvate the protein. YidC was inserted into the lipid bilayer, solvated, and ionized using the membrane builder on CHARMM-GUI. ^47^ In these MD studies, we used palmitoyloleoyl Phosphatidylethanolamine (POPE) lipids to build a lipid bilayer. A membrane layer surface of 110 Å × 110 Å was built along the XY plane. The protein lipid-assembly was solvated in water with 25 Å thick layers of water on top and bottom. 0.15 M of Na^+^ and Cl^*−*^ ions were added to the solution with a slight modification in the number of ions to neutralize the system. The solvated system contained ≈142000 atoms. Before the equilibrium simulation, the structure was energy minimized using the conjugate gradient algorithm.^48^ Following that, we used the standard CHARMM-GUI^47^ protocol to progressively relax the systems using restrained MD simulations. In the NPT ensemble at 310 K, 550 ns of equilibrium MD simulations were performed under periodic boundary conditions for each system. In the simulations, a Langevin integrator with a damping coefficient of *γ* =0.5 ps^*−*1^ and 1 atm pressure was maintained using the Nose-Hoover Langevin piston method ^49,50^ in the simulations.

**Fig. 2.**
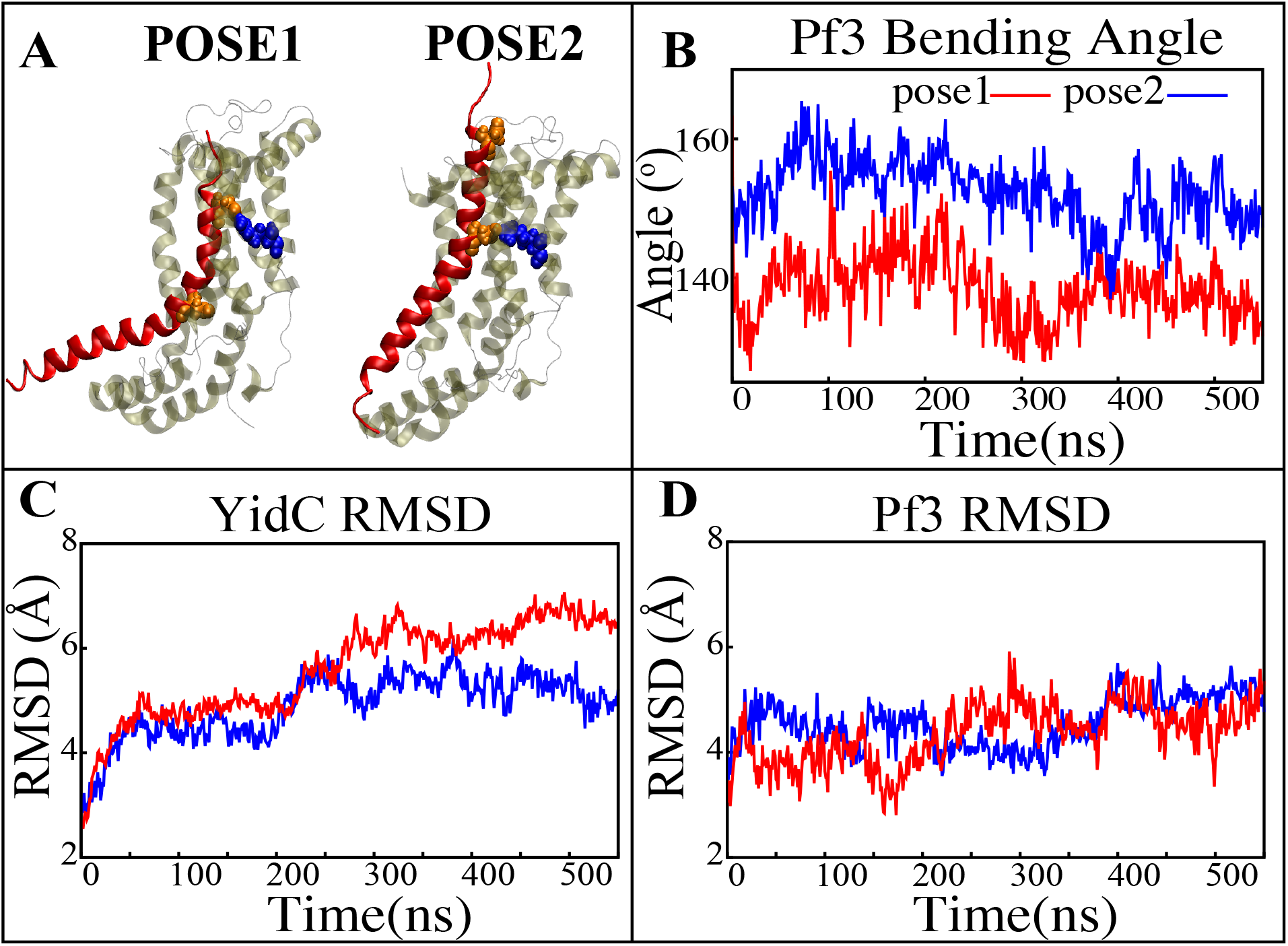
Structural stability evaluation of YidC and Pf3 in the insertion process. (A) pose1 & pose2 docking models based on the YidC Pf3 coat insertion process, as described previously. (B) The bending angle of the Pf3 helix in pose1 (red) and pose2 (blue) models shown as a function of time. (C & D) Root mean square deviation of the YidC & Pf3 coat in pose1 (red) and pose2 (blue) models. Based on RMSD results, we have observed that in pose2 stage the YidC is significantly more stable compared to the pose1 state.

Trajectories were visualized and analyzed using VMD software.^51^ Salt bridge interaction analysis was conducted via VMD plugins. For salt bridge analysis, the cut-off distance was set at 4 Å and the distance between the oxygen atoms of the acidic residues and the nitrogen atoms of basic residues was calculated. The interhelical angles were calculated as the angle between the third principal axes of the corresponding helices.^52–54^ The TM helices and other sub-domains were defined for analysis as follows: TM1a (58*−*78), TM1b (79*−*104), TM2 (134*−*155), TM3 (175*−*190), TM4 (219*−*233), TM5 (233*−*258), C1 region (84*−*133), C2 loop (195*−*216), and modified C-terminal region (256*−*272) respectively. The number of contacts within 3 Å of selection was measured for contact analysis. We counted the number of water molecules within 5 Å of R72 for water analysis. For the Pf3 bending angle, we chose two pairs of residues selection for the top (ASP7-ASP18) and bottom (ASP18-LEU29) regions of Pf3 respectively and measured their third principal axes, denoted by *v*_1_ and *v*_2_, respectively. The angle between the two vectors was calculated as 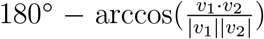. Principal component analysis (PCA) was performed on each trajectory using PRODY^55^ software. Only *C*_*α*_ atoms of the peptide were considered in the PCA calculations of both docking simulations.

The combination of equilibrium and non-equilibrium MD simulations has proven effective for investigating biological challenges. ^12,56–64^ In this work, the YidC independent insertion mechanism was studied using non-equilibrium targeted MD (TMD) as implemented within the colvars module of NAMD.^65^ TMD simulation was performed on the final conformation of the pose 1 system obtained from the 550 ns equilibrium trajectory in order to transfer the Pf3 peptide to the periplasmic side of the membrane, as seen in pose 2. The RMSD collective variable was used in the TMD simulation. As a collective variable, we used the RMSD of Pf3 Coat’s backbone atoms from the last frame of pose2’s equilibrium simulation trajectory. TMD simulation was run for 100 ns with a force constant of 44 *kcal/mol/*Å^2^. To ensure conformational accuracy, the final frame of the targeted MD simulation was equilibrated for 20 ns without any restraints.

## Results and Discussion

### YidC Undergoes Major Conformational Changes in Sec-independent Insertion Process

A protein must undergo various local and global conformational changes in a mechanism. A set of approximate docking models (Fig. 2A) of YidC and Pf3 were developed to represent the insertion process. MD simulations of these docking models were performed to investigate various conformational characteristics to see how YidC and Pf3 altered conformational properties during the insertion process. Several measures or quantities linked to local and global conformational properties were evaluated and monitored for the two conformational poses developed in this study. The C*α* root mean square deviation (RMSD) of the systems were evaluated independently of the framework to test the impact of the Pf3 protein on YidC’s global structure. It was found that the Pf3 coat had a relatively comparable RMSD in the two models simulated in this study (Fig. 2D). However, the YidC protein fluctuated more in pose 1 (Fig. 2C) than in pose 2 (Fig. 2C). At the start of a process or mechanism, a protein is anticipated to undergo substantial conformational changes. The fact that YidC’s RMSD in pose 1 (Fig. 2C) is 2 Å greater than that in pose 2 (Fig. 2C) suggests that YidC goes through significant conformational changes at the start of the process. This demonstrates that the effect of Pf3 insertion into the membrane differs depending on the stage of the process. Although we see comparable RMSD of Pf3 in both poses, the Pf3 bending angle analysis (Fig. 2A) more clearly suggests a structural difference between the two states in support of our hypothesis. The bending angle indicates that Pf3 has a lower bend at the start of the insertion process and changes its conformation inside the YidC groove (Fig. 2A) as it progresses deeper into the groove. This brings us to the conclusion, that Pf3 undergoes major conformational changes to adapt to the YidC groove environment during the insertion process. In the next investigation, additional local components of YidC were rigorously investigated to elucidate more details of the insertion process.

### Widening of the Transmembrane Domain is Essential for Incorporation of Proteins in Membrane During the Insertion Process

Previous studies have revealed that the YidC transmembrane (TM) region is crucial for membrane protein insertion into the lipid bilayer.^66,67^ The helical angle between each TM pair was determined in this study. In the pose2 docking simulation, the transmembrane helices (TM1a, TM2, TM3, TM4, and TM5) seem more slanted than in the pose1 docking simulation, which has a difference in the angle of over 10 degrees (Fig. 3B-G). This implies that the central TM groove of YidC is substantially enlarged during the insertion of Pf3 into the membrane bilayer. The critical helices TM1a (Fig. 3E,F,&G) and TM2 (Fig. 3B,C,&D) undergo significant modifications following peptide insertion because they are stretched onto the cytoplasmic side of the membrane, which is the entrance point of Pf3. Based on this, one may assume that throughout the insertion process, YidC experiences a gradual and tranquil conformational shift, organically adjusting to the incoming peptide. In this scenario, Yidc progressively expands its transmembrane groove to make room for the incoming peptide and then returns to a normal state once the peptide is fully incorporated into the membrane.

**Fig. 3.**
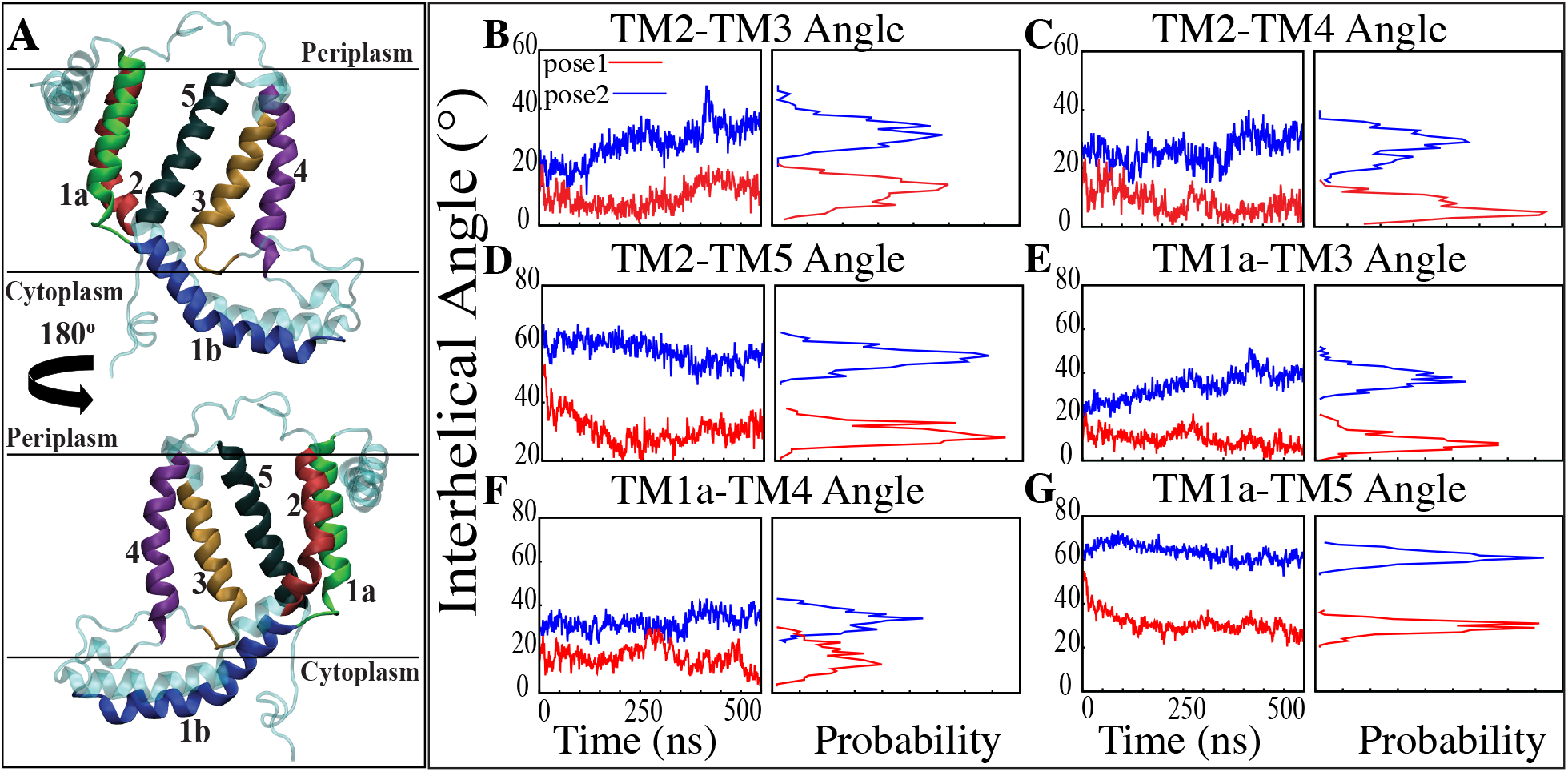
Inter-helical angles between trans-membrane helices of YidC in both docking model simulations. (A) Graphical representation of YidC protein on periplasmic, cytoplasmic and transmembrane regions labeled with helical numbers in the transmembrane region. (B-D) Overall inter-helical angle between transmembrane helix 2 and other helices of the protein in pose 1 (red) and pose 2 (blue) simulations. (E-G) Overall inter-helical angle between transmembrane helix 1a and other helices of the protein in pose 1 and pose 2 simulations. Also, the probability density distribution is shown for all graphs.

The interactions of the membrane and YidC with the Pf3 coat were studied to learn more about the insertion process. The number of interactions of Pf3 with YidC and the membrane within 3 Å were taken into account for the interaction analysis (Fig. 4A&B). The contact of the Pf3 coat with the membrane determines its position in the bilayer system; the Pf3 coat positioned inside YidC’s hydrophilic groove has a greater lipid interaction distribution than the Pf3 coat positioned just outside the groove area (Fig. 4B). Because the Pf3 coat is now ready to be incorporated into the membrane, Pf3 has a high degree of contact with the membrane in pose2. The divergence of Pf3 lipid interactions supports the suggested mechanistic models modeled for the YidC independent insertion pathway in this study. In addition to lipid interactions, the interactions between YidC and Pf3 are also important in this process. The distribution of such interactions should confirm the outcomes of the lipid interactions. Because of the greater distribution of lipid contacts in the pose2 model, Pf3 decreases its interaction with the YidC protein by shifting closer to the lipid bilayer (Fig. 4B). Whereas at the pose1 stage of the insertion process, the association of YidC and signal protein should be significantly strengthened before establishing the peptide in YidC’s hydrophilic groove. This would explain the YidC higher interactions with the Pf3 coats that were observed in the pose1 model (Fig. 4A).

**Fig. 4.**
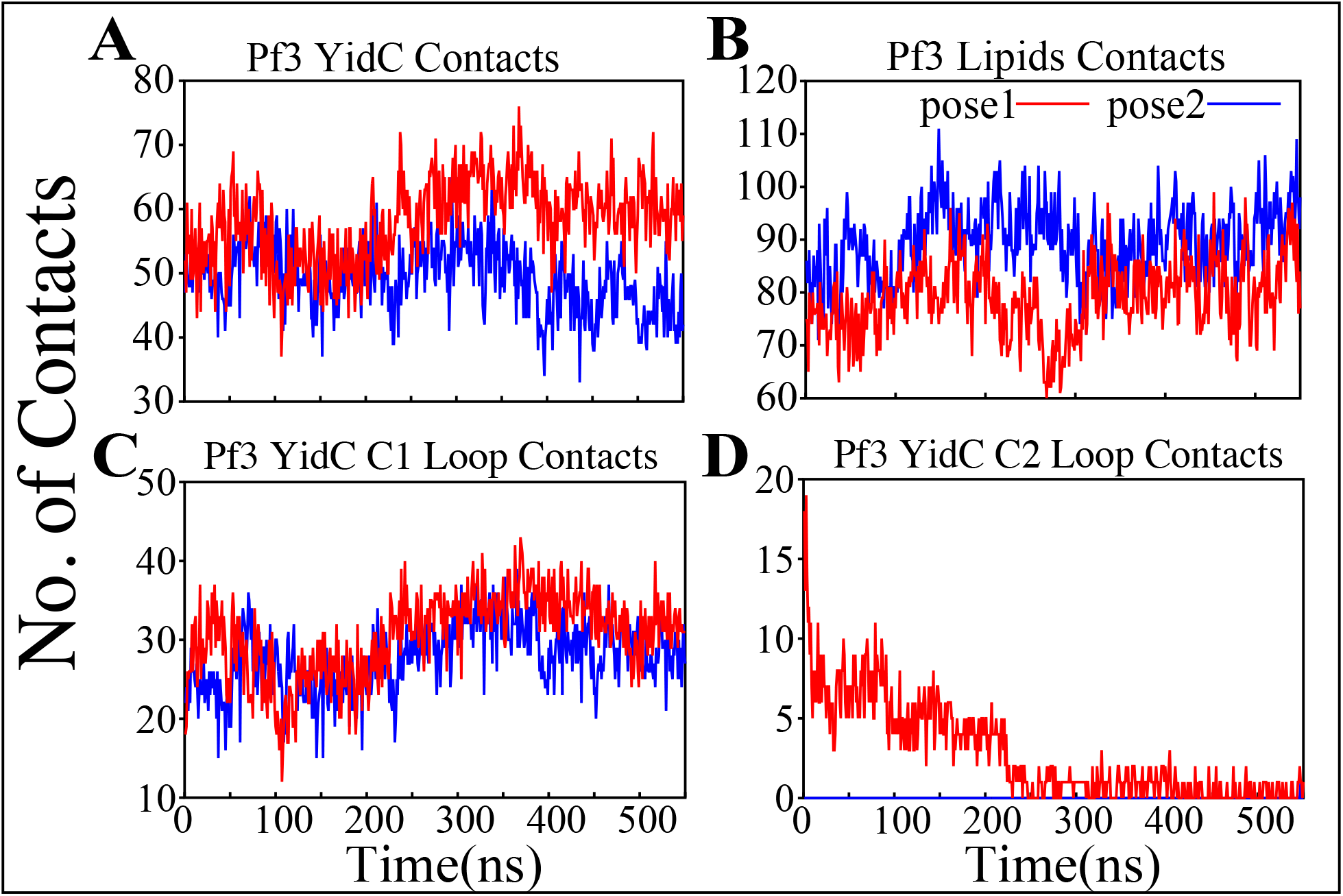
Pf3 overall interaction with YidC and POPE lipid tails in both the docking model simulations. (A & B) Respective number of YidC and lipid interactions with the Pf3 in pose1 (red) and pose2 (blue), shown as a function of time. (C & D) Number of contacts between Pf3 coat and the C1 and C2 loops of YidC, shown as a function of time.

### Interaction of C1 and C2 Loops with Pf3 Stabilizes the Insertion Process

Cytoplasmic loops C1 and C2 (Fig. 1A) are important components in YidC’s independent insertion mechanism. Previous studies on YidC with or without the C2 loop found that its presence stabilizes YidC in the membrane.^68^ In both the pose1 and pose2 models, the YidC cytoplasmic loop C1 interacts with the Pf3 coat protein. This interaction aids in the stability of the Pf3 coat inside YidC’s hydrophilic groove. Furthermore, these loops establish a strong interaction to retain the signal proteins inside YidC’s U-shaped groove (Fig. 4C&D). At the beginning of the insertion mechanism, the cytoplasmic C1 loop, which is deeply expanded into the cytoplasmic side, creates extremely strong contacts with the Pf3 protein (Fig. 4C). The interactions between the C1 loop and the peptide are critical for keeping the peptide under control during the insertion process. According to our contact analysis results, the C2 loop contacts (Fig. 4D) are formed only in the pose1 model, since it is the starting point of the insertion process, and a high number of protein interactions are necessary to stabilize such a long peptide. As the process progresses, the C2 loop loses its interactions (Fig. 4D) with the incoming peptide once the peptide is settled inside the U-shaped groove of YidC, as seen in pose2. Thus far, we have shown that YidC undergoes significant global and local conformational changes, such as TM domain expansion, and its interactions with Pf3, specifically through contacts in the cytoplasmic loop region. All the findings presented above confirm the major hypothesis of the study on YidC conformational changes throughout the independent insertion process.

Principal component analysis (PCA) was used to identify the key differences between the pose1 and pose2 models. Pose1 and pose2 systems were clearly differentiated by projections onto principal components (PCs) 1 and 2. Only YidC C_*α*_ atoms are considered in this study.

PC1 and PC2 contributed 64.4 and 47.9 percent of the total variance, respectively (Fig. 5A). As expected, the structural analysis of pose1 and pose2 models contradicts each other in PC1 and PC2, which is logical given the significant conformational differences (Fig. 5A) observed previously. The Pf3 coat, on the other hand, has clustered similarly along PC1 (Fig. 5B). However, the fluctuation of Pf3 is different around PC2 (Fig. 5B), which may be the result of a shift in Pf3 interactions and conformational changes. To demonstrate this visually, square displacements of PC residues were projected onto the structure, as seen in Fig. 5C-D. Overall, the major finding of the PC analysis was that the behavior of the pose1 and pose2 proteins differed considerably. This confirms our previous notion that YidC conformational dynamics play an important role in the insertion process. The PCA results are consistent with the early evidence for global and local structural changes.

**Fig. 5.**
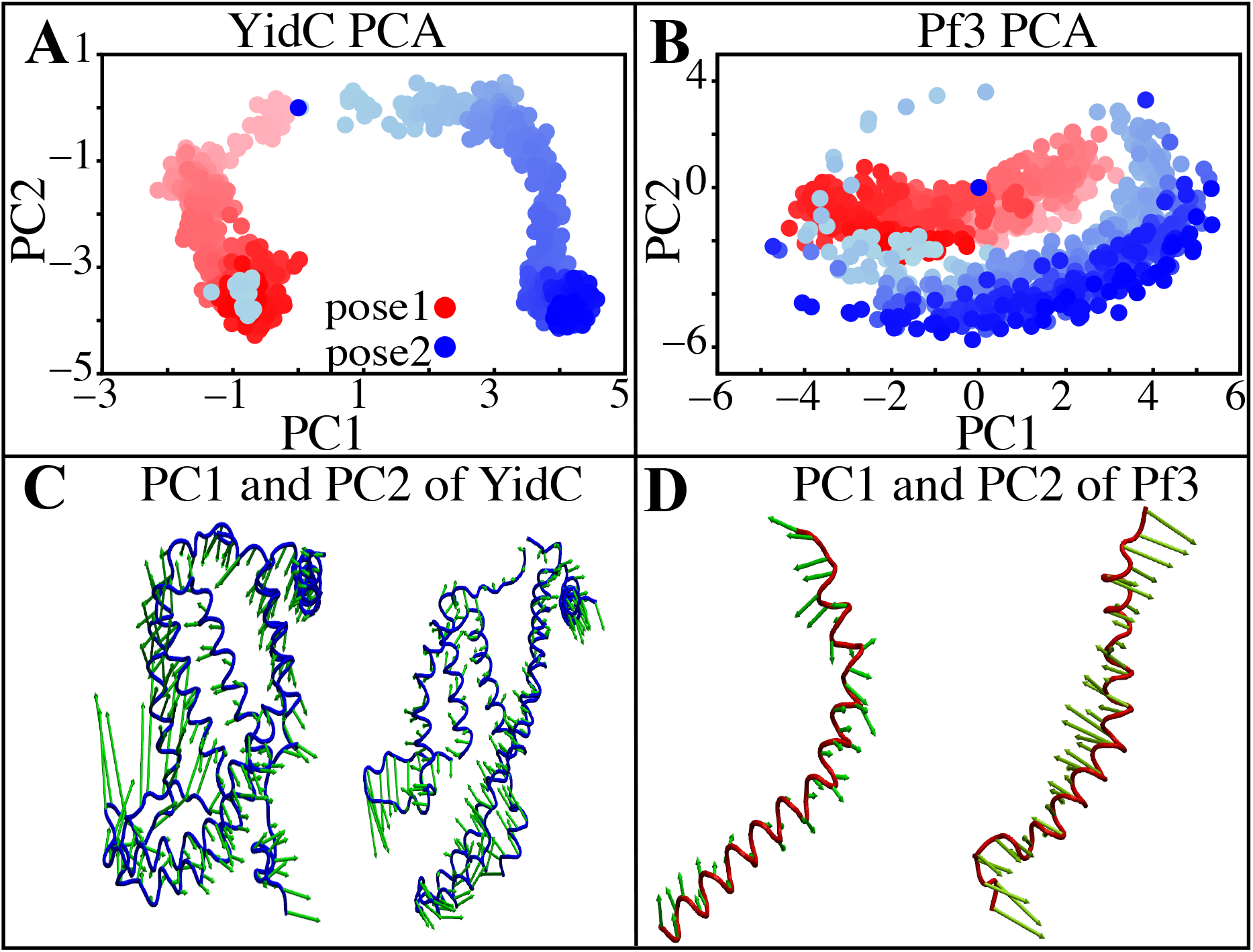
Principal component projections along PCs 1 and 2. (A & B) PC1 vs PC2 for pose1(red) and pose2(blue) models of YidC & Pf3 coat. (C) Structural variation in PC1 and PC2 of YidC, respectively. (D) Structural variations in PC1 and PC2 of Pf3, respectively. The bidirectional arrow shows the direction of the fluctuation of the structure and the length of the arrow reflects the magnitude of the fluctuation. The color shading in the picture indicates a timeline, with light and dark shades representing the beginning and end of the simulation, respectively.

### YidC’s Hydrophilic Groove Hydration and Dehydration are Critical in the Insertion Mechanism

YidC has a U-shaped hydrophilic groove that is closed on the periplasmic side but exposed to the cytoplasmic side of the membrane bilayer. To examine the water content of the groove within helices TM1-TM5 (Fig. 6A), the number of water molecules inside the groove region of the YidC protein was measured and plotted over the simulation time. The water analysis results reveal that the number of water molecules within the groove region is higher in pose 1, which is considered the starting state of the insertion process. Whereas in the docking model pose2, the water content is close to zero throughout the simulation (Fig. 6B). This confirms the previous hypothesis that a water slide motion is important in the initial positioning of the Pf3 coat protein.^38,39^ The peptide enters the YidC groove via the cytoplasmic side of the membrane bilayer; the central TM helices are then widened to form a water slide^8^ and the YidC groove region is filled with water to provide a smooth sliding motion for the entering protein. As Pf3 progresses through the insertion processes, the cytoplasmic groove of YidC becomes more compact and water molecules are pushed out of the TM groove. These two factors combine to cause a hydrophobic shift in the region, making it more susceptible to membrane insertion. This results in an overall water slide motion for an insertion of protein inside the YidC’s TM helical region.

**Fig. 6.**
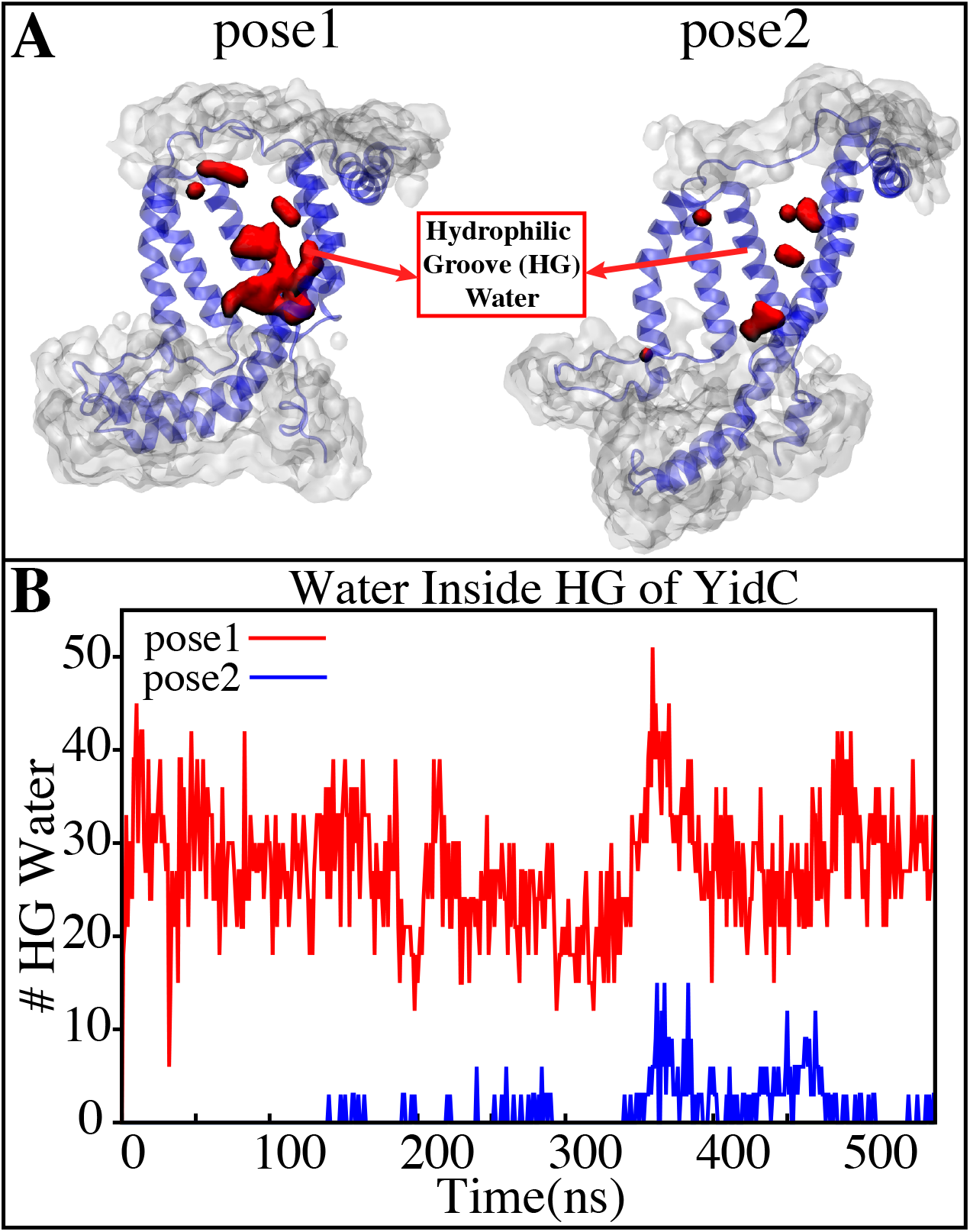
(A) Graphical representation of the docking models showing the average water molecule count in the hydrophilic groove (HG) region of YidC. (B) Number of water molecules inside the hydrophilic groove (HG) region of YidC in docking poses 1 (red) & pose 2 (blue). Pf3 entry into the TM area is aided by the water in the groove, which creates a sliding action.

### The Saltbridge Interaction of Pf3 with YidC R72 in the Hydrophilic Groove is a Significant Event in the Insertion Process

The YidC residue Arginine 72 (R72) is in the core cavity of the YidC transmembrane region and forms a salt-bridge with incoming protein chains. It has been suggested that before translocation, a YidC protein’s hydrophilic groove is forced into the hydrophobic cavity, implying that peptides may reach R72 for bond formation.^69^ According to salt bridge analysis results, R72 is available for interacting with the incoming Pf3 coat protein. During the insertion process, the R72 residue of YidC forms a stable salt-bridge with D7 and D18 of Pf3 in the pose1 and pose2 simulations, respectively (Fig. 7). During the first phase of YidC insertion, the salt-bridge interaction between YidC’s R72 and Pf3’s D7 stabilizes the Pf3 coat in the TM helical groove as soon as it enters the TM groove. As the Pf3 coat moves towards the periplasmic side of the protein, salt-bridge residue interactions with the Pf3 coat are sequentially moved from D7 to D18 (Fig. 8A).

**Fig. 7.**
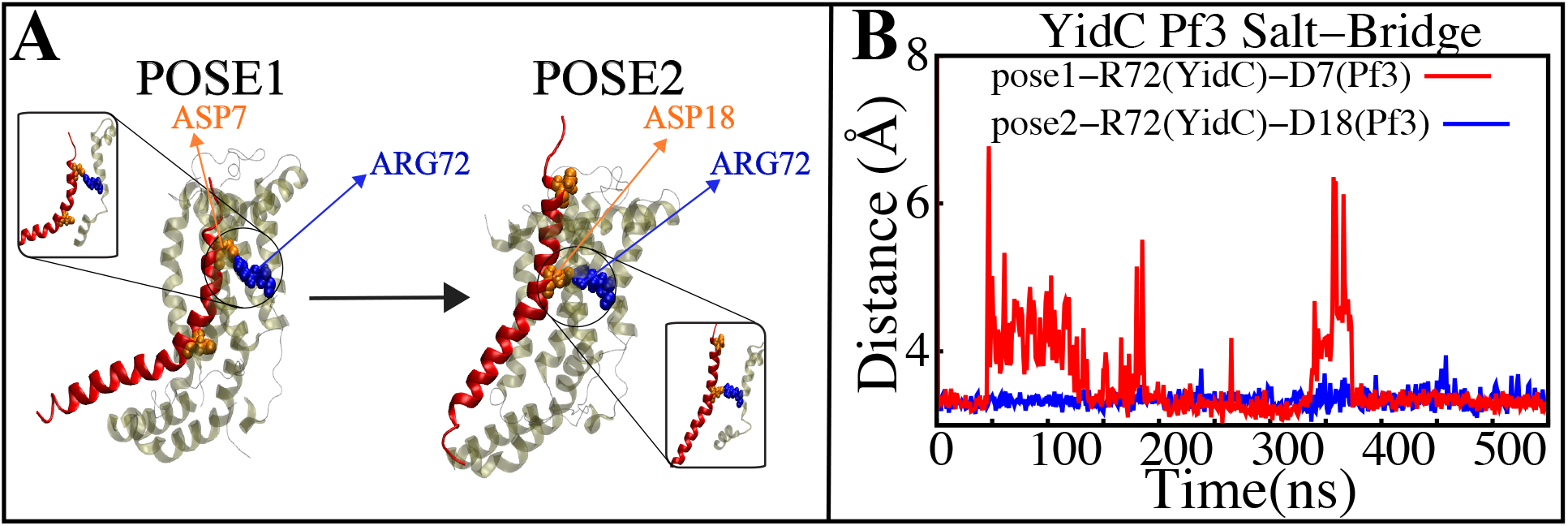
Salt−bridge connectivity of R72 (YidC) located in the groove. (A) Graphical representation of significant salt-bridge interactions between the Pf3 coat and YidC involved in the insertion process. (B) Distance between salt-bridge Arg72(YidC)−Asp18(Pf3 coat) and Arg72(YidC)−Asp7(Pf3 coat) (labeled with blue and red lines, respectively) in YidC and Pf3 coat docking models.

**Fig. 8.**
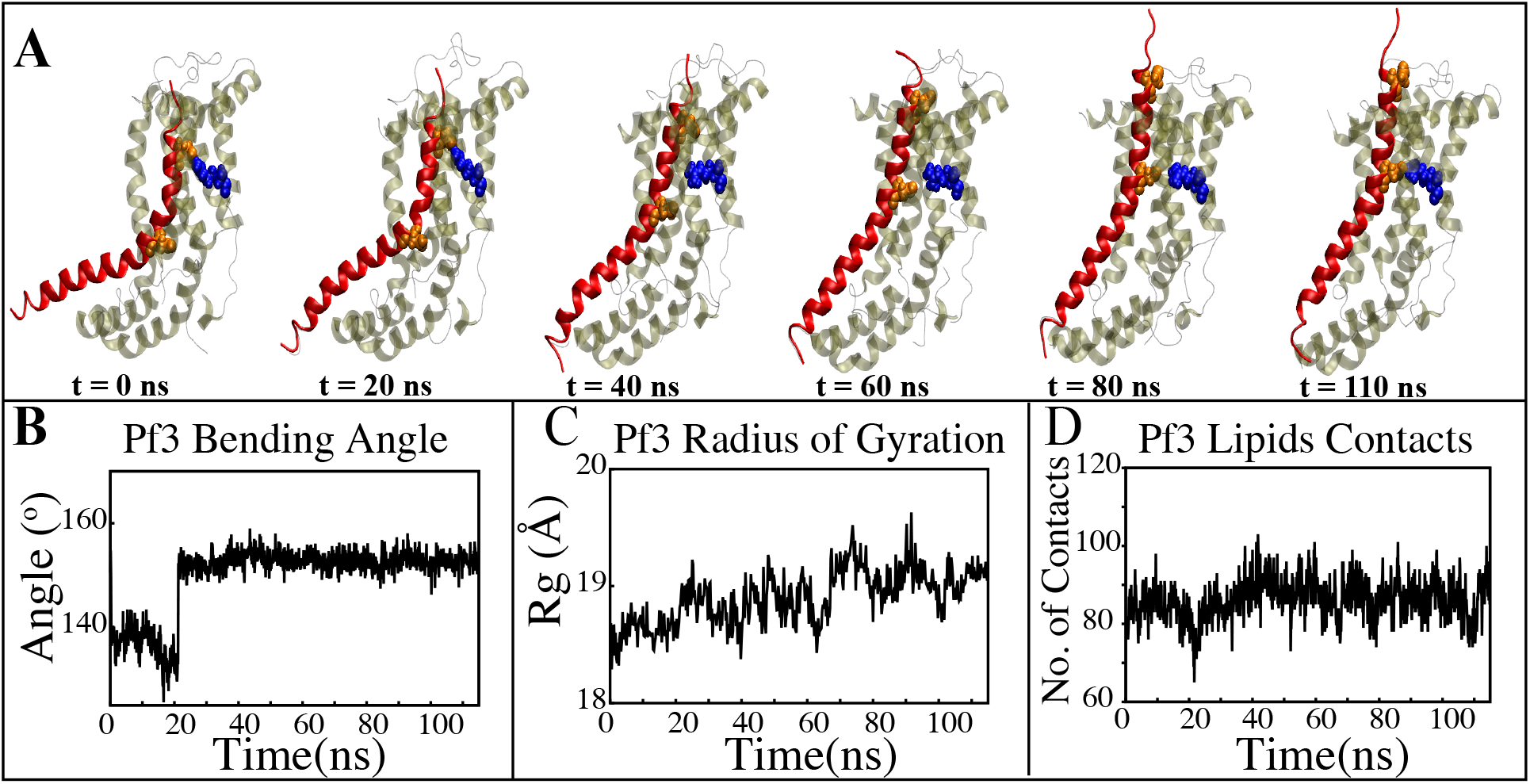
Characterizing the insertion process using targeted MD simulations. (A) Graphical representation of a series of targeted MD snapshots taken at different stages of the simulation. (B) Bending angle analysis of the Pf3 helix. (C) Radius of gyration of Pf3 peptide in the insertion process. (D) Interactions of Pf3 with lipid tails in the NE simulation process.

### Non-Equilibrium Simulation of YidC’s Sec-independent Mechanism of Pf3 Coat Insertion in the Membrane Bilayer

The insertion process was further investigated using the above-mentioned non-equilibrium (NE) simulation (Fig. 8A) approach. Many of the key factors discussed above, such as Pf3 bending angle (Fig. 8B), radius of gyration (Fig. 8C), Pf3 lipid interactions (Fig. 8D), the presence of water in the groove (Fig. 9C), and Pf3 contacts with YidC (Fig. 9D), are evaluated for the NE simulation trajectory. Our NE simulation results are totally in agreement with results produced in equilibrium simulations. The bending of Pf3 is observed in the NE simulations, where Pf3 has gone from a lower to a greater bending angle (Fig. 8B) to adapt to the groove environment. The radius of gyration analysis also confirms our hypothesis about Pf3 conformational changes during the insertion process (Fig. 8C). The increase and decrease in the amount of water inside the groove significantly supports the hydration and dehydration hypothesis (Fig. 9C). Robust interactions of Pf3 with lipid tails (Fig. 8D) play a significant role in the insertion process. As previously stated, YidC loses connections with the Pf3 coat as the insertion process progresses, as seen in the NE simulations, where the number of YidC-Pf3 contacts decreases during the targeted MD simulation (Fig. 9D). During the insertion of Pf3 inside the membrane, YidC undergoes significant conformational changes, which we observed previously in our analysis. As expected, Yidc underwent substantial conformational changes from the beginning to the completion of the insertion process as indicated by the overall RMSD (Fig. 9A) and radius of gyration (Fig. 9B) analyses. Overall, based on equilibrium and NE simulations, the following mechanism for YidC’s Sec-independent insertion mechanism is proposed in this study: The incoming Pf3 coat protein first interacts with the cytoplasmic loops and gradually moves into the hydrophilic groove located in the transmembrane region, forming a salt bridge with R72. The negatively charged D7 residue of the Pf3 coat protein forms a salt bridge with the positively charged R72. The formation of this salt bridge is critical to the insertion mechanism. The Pf3 coat protein then migrates towards the periplasmic side of the membrane, assisted by a force generated by pushing water out of the YidC hydrophilic groove. A salt bridge is then formed between D18 of Pf3 and R72 of YidC to stabilize the position of Pf3 in the membrane. Later in the process, Pf3 moves into the membrane. It leaves the hydrophobic TM region and moves to the lipid bilayer.

**Fig. 9.**
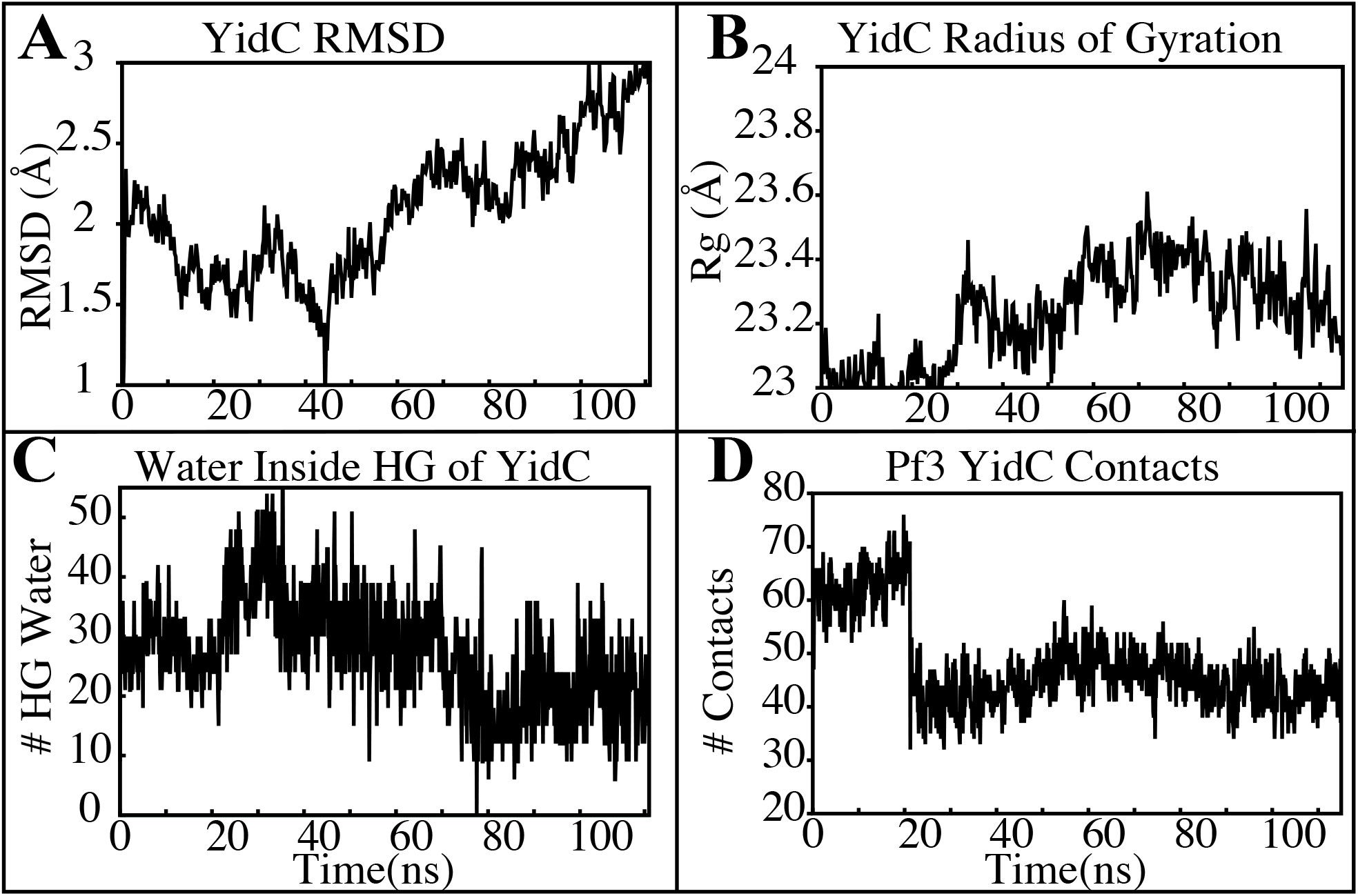
YidC conformational changes observed in the targeted MD simulations. (A & B) Root mean square deviation and radius of gyration of YidC in the insertion process of Pf3. (C) The water content in the hydrophilic groove of the protein during a 100ns NE simulation followed by a 15 ns equilibrium simulation. (D) Interaction of YidC with Pf3 in the NE simulation process.

## Conclusions

Based on our equilibrium and non-equilibrium MD simulation results, YidC must undergo major conformational changes during the secY-independent insertion process. The incoming Pf3 coat protein would first come into contact with the cytoplasmic loops and then penetrate into the hydrophilic groove, forming a salt bridge with R72. The YidC loops on the cytoplasmic side of the bilayer are critical for moving Pf3 into YidC’s hydrophilic groove. At first, these cytoplasmic loops make contact with the Pf3 coat. The negatively charged D7 residue of Pf3 interacts with the positively charged R72 of YidC to form a stable salt bridge. The formation of this salt bridge is crucial in the insertion process to stabilize the Pf3 in the YidC’s TM groove. The Pf3 coat protein then travels towards the periplasmic side of the membrane, helped by the water slide force. The interactions with the membrane also aid in the passage of the protein towards the periplasmic side, which is also supported by the salt bridge between D18 of Pf3 and R72 of YidC; this combination stabilizes the position of Pf3 in the membrane. The protein then moves into the membrane through the water-filled cleft. Finally, when the Pf3 coat is completely inside the YidC’s hydrophilic groove, it will come into contact with lipid tails. This will be helped by the hydration of the groove, which will push the Pf3 coat into the bilayer.

## Acknowledgement

This research is supported by National Science Foundation grant CHE 1945465 and the Arkansas Biosciences Institute. This research is part of the Blue Waters sustained petascale computing project, which is supported by the National Science Foundation (awards OCI-0725070 and ACI-1238993) and the state of Illinois. This work also used the Extreme Science and Engineering Discovery Environment (allocation MCB150129), which is supported by National Science Foundation grant number ACI-1548562. This research is also supported by the Arkansas High-Performance Computing Center, which is funded through multiple National Science Foundation grants and the Arkansas Economic Development Commission.

## References

1. Rapoport, T. A. Protein translocation across the eukaryotic endoplasmic reticulum and bacterial plasma membranes. Nature 2007, 450, 663–669.

2. Krogh, A.; Larsson, B.; Von Heijne, G.; Sonnhammer, E. L. Predicting transmembrane protein topology with a hidden Markov model: Application to complete genomes. Journal of Molecular Biology 2001, 305, 567–580.

3. Dalbey, R. E.; Kuhn, A. Membrane Insertases Are Present in All Three Domains of Life. Structure 2015, 23, 1559–1560.

4. McDowell, M. A.; Heimes, M.; Sinning, I. Structural and molecular mechanisms for membrane protein biogenesis by the Oxa1 superfamily. Nature Structural & Molecular Biology 2021, 28, 234–239.

5. Jiang, F.; Chen, M.; Yi, L.; De Gier, J. W.; Kuhn, A.; Dalbey, R. E. Defining the regions of Escherichia coli YidC that contribute to activity. Journal of Biological Chemistry 2003, 278, 48965–48972.

6. Nass, K. J.; Ilie, I. M.; Saller, M. J.; Driessen, A. J. M.; Caflisch, A.; Kammerer, R. A.; Li, X. The role of the N-terminal amphipathic helix in bacterial YidC: Insights from functional studies, the crystal structure and molecular dynamics simulations. Biochimica et Biophysica Acta (BBA) - Biomembranes 2022, 1864, 183825.

7. Güngör, B.; Flohr, T.; Garg, S. G.; Herrmann, J. M. The ER membrane complex (EMC) can functionally replace the Oxa1 insertase in mitochondria. PLOS Biology 2022, 20, e3001380.

8. Dalbey, R. E.; Kuhn, A.; Zhu, L.; Kiefer, D. The membrane insertase YidC. Biochimica et Biophysica Acta - Molecular Cell Research 2014, 1843, 1489–1496.

9. Dalbey, R. E.; Chen, M. Sec-translocase mediated membrane protein biogenesis. Biochimica et Biophysica Acta - Molecular Cell Research 2004, 1694, 37–53.

10. Borowska, M. T.; Dominik, P. K.; Anghel, S. A.; Kossiakoff, A. A.; Keenan, R. J. A YidC-like Protein in the Archaeal Plasma Membrane. Structure 2015, 23, 1715–1724.

11. Kuhn, A.; Kiefer, D. Membrane protein insertase YidC in bacteria and archaea. Molecular Microbiology 2017, 103, 590–594.

12. Chen, H.; Ogden, D.; Pant, S.; Cai, W.; Tajkhorshid, E.; Moradi, M.; Roux, B.; Chipot, C. A Companion Guide to the String Method with Swarms of Trajectories: Characterization, Performance, and Pitfalls. Journal of Chemical Theory and Computation 2022, 18, 1406–1422.

13. Scotti, P. A.; Urbanus, M. L.; Brunner, J.; De Gier, J. W. L.; Von Heijne, G.; Van Der Does, C.; Driessen, A. J.; Oudega, B.; Luirink, J. YidC, the Escherichia coli homologue of mitochondrial Oxa1p, is a component of the Sec translocase. EMBO Journal 2000, 19, 542–549.

14. Facey, S. J.; Kuhn, A. Membrane integration of E. coli model membrane proteins. Biochimica et Biophysica Acta - Molecular Cell Research 2004, 1694, 55–66.

15. Lewis, N. E.; Brady, L. J. Breaking the bacterial protein targeting and translocation model: Oral organisms as a case in point. Molecular Oral Microbiology 2015, 30, 186– 197.

16. Samuelson, J. C.; Chen, M.; Jiang, F.; Möller, I.; Wiedmann, M.; Kuhn, A.; Phillips, G. J.; Dalbey, R. E. YidC mediates membrane protein insertion in bacteria. Nature 2000, 406, 637–641.

17. Kiefer, D.; Kuhn, A. YidC-mediated membrane insertion. FEMS Microbiology Letters 2018, 365.

18. Laskowski, P. R.; Pluhackova, K.; Haase, M.; Lang, B. M.; Nagler, G.; Kuhn, A.; Müller, D. J. Monitoring the binding and insertion of a single transmembrane protein by an insertase. Nature Communications 2021, 12, 7082.

19. Lewis, A. J. O.; Hegde, R. S. A unified evolutionary origin for the ubiquitous protein transporters SecY and YidC. BMC Biology 2021, 19, 266.

20. Nagamori, S.; Smirnova, I. N.; Kaback, H. R. Role of YidC in folding of polytopic membrane proteins. Journal of Cell Biology 2004, 165, 53–62.

21. Serdiuk, T.; Balasubramaniam, D.; Sugihara, J.; Mari, S. A.; Kaback, H. R.; Müller, D. J. YidC assists the stepwise and stochastic folding of membrane proteins. Nature Chemical Biology 2016, 12, 911–917.

22. Kol, S.; Turrell, B. R.; De Keyzer, J.; Van Der Laan, M.; Nouwen, N.; Driessen, A. J. YidC-mediated membrane insertion of assembly mutants of subunit c of the F1F0 ATPase. Journal of Biological Chemistry 2006, 281, 29762–29768.

23. Van der Laan, M.; Urbanus, M. L.; Ten Hagen-Jongman, C. M.; Nouwen, N.; Oudega, B.; Harms, N.; Driessen, A. J.; Luirink, J. A conserved function of YidC in the biogenesis of respiratory chain complexes. Proceedings of the National Academy of Sciences of the United States of America 2003, 100, 5801–5806.

24. Yi, L.; Dalbey, R. E. Oxa1/Alb3/YidC system for insertion of membrane proteins in mitochondria, chloroplasts and bacteria. Molecular Membrane Biology 2005, 22, 101– 111.

25. Van Bloois, E.; Jan Haan, G.; De Gier, J. W.; Oudega, B.; Luirink, J. F1F0 ATP synthase subunit c is targeted by the SRP to YidC in the E. coli inner membrane. FEBS Letters 2004, 576, 97–100.

26. Endo, Y.; Shimizu, Y.; Nishikawa, H.; Sawasato, K.; Nishiyama, K.-i. Interplay between MPIase, YidC, and PMF during Sec-independent insertion of membrane proteins. Life Science Alliance 2022, 5.

27. Xin, Y.; Zhao, Y.; Zheng, J.; Zhou, H.; Zhang, X. C.; Tian, C.; Huang, Y. Structure of YidC from Thermotoga maritima and its implications for YidC-mediated membrane protein insertion. FASEB Journal 2018, 32, 2411–2421.

28. Yuan, J.; Phillips, G. J.; Dalbey, R. E. Isolation of cold-sensitive yidC mutants provides insights into the substrate profile of the YidC insertase and the importance of transmembrane 3 in YidC function. Journal of Bacteriology 2007, 189, 8961–8972.

29. Klenner, C.; Kuhn, A. Dynamic disulfide scanning of the membrane-inserting Pf3 coat protein reveals multiple YidC substrate contacts. Journal of Biological Chemistry 2012, 287, 3769–3776.

30. Kol, S.; Nouwen, N.; Driessen, A. J. Mechanisms of YidC-mediated insertion and assembly of multimeric membrane protein complexes. Journal of Biological Chemistry 2008, 283, 31269–31273.

31. Ernst, S.; Schönbauer, A. K.; Bär, G.; Börsch, M.; Kuhn, A. YidC-driven membrane insertion of single fluorescent Pf3 coat proteins. Journal of Molecular Biology 2011, 412, 165–175.

32. Funes, S.; Kauff, F.; Van Der Sluis, E. O.; Ott, M.; Herrmann, J. M. Evolution of YidC/Oxa1/Alb3 insertases: Three independent gene duplications followed by functional specialization in bacteria, mitochondria and chloroplasts. Biological Chemistry 2011, 392, 13–19.

33. Funes, S.; Hasona, A.; Bauerschmitt, H.; Grubbauer, C.; Kauff, F.; Collins, R.; Crowley, P. J.; Palmer, S. R.; Brady, L. J.; Herrmann, J. M. Independent gene duplications of the YidC/Oxa/Alb3 family enabled a specialized cotranslational function. Proceedings of the National Academy of Sciences of the United States of America 2009, 106, 6656–6661.

34. Kohler, R.; Boehringer, D.; Greber, B.; Bingel-Erlenmeyer, R.; Collinson, I.; Schaffitzel, C.; Ban, N. YidC and Oxa1 Form Dimeric Insertion Pores on the Translating Ribosome. Molecular Cell 2009, 34, 344–353.

35. Chen, Y.; Capponi, S.; Zhu, L.; Gellenbeck, P.; Freites, J. A.; White, S. H.; Dalbey, R. E. YidC Insertase of Escherichia coli: Water Accessibility and Membrane Shaping. Structure 2017, 25, 1403–1414.e3.

36. Ito, S.; D’Alessio, A. C.; Taranova, O. V.; Hong, K.; Sowers, L. C.; Zhang, Y. Role of Tet proteins in 5mC to 5hmC conversion, ES-cell self-renewal and inner cell mass specification. Nature 2010, 466, 1129–1133.

37. Kedrov, A.; Wickles, S.; Crevenna, A. H.; van der Sluis, E. O.; Buschauer, R.; Berninghausen, O.; Lamb, D. C.; Beckmann, R. Structural Dynamics of the YidC:Ribosome Complex during Membrane Protein Biogenesis. Cell Reports 2016, 17, 2943–2954.

38. Wickles, S.; Singharoy, A.; Andreani, J.; Seemayer, S.; Bischoff, L.; Berninghausen, O.; Soeding, J.; Schulten, K.; van der Sluis, E. O.; Beckmann, R. A structural model of the active ribosome-bound membrane protein insertase YidC. eLife 2014, 3, 1–17.

39. Ito, K.; Shimokawa-Chiba, N.; Chiba, S. Sec translocon has an insertase-like function in addition to polypeptide conduction through the channel. F1000Research 2019, 8, F1000 Faculty Rev–2126.

40. Fujihashi, M.; Mito, K.; Pai, E. F.; Miki, K. Atomic resolution structure of the orotidine 5’-monophosphate decarboxylase product complex combined with surface plasmon resonance analysis: implications for the catalytic mechanism. The Journal of biological chemistry 2013, 288, 9011–9016.

41. Molecular Operating Environment (MOE),. Chemical Computing Group Inc. Molecular Operating Environment (MOE); Chemical Computing Group Inc. 1010 Sherbooke St. West, Suite# 910: Montreal, QC, Canada,. Molecular Operating Environment (MOE), 2013.08; Chemical Computing Group Inc., 1010 Sherbooke St. West, Suite #910, Montreal, QC, Canada, H3A 2R7, 2013. 2015,

42. Tsukazaki, T. Structural Basis of the Sec Translocon and YidC Revealed Through X-ray Crystallography. The Protein Journal 2019, 38, 249–261.

43. Phillips, J. C.; Braun, R.; Wang, W.; Gumbart, J.; Tajkhorshid, E.; Villa, E.; Chipot, C.; Skeel, R. D.; Kalé, L.; Schulten, K. Scalable molecular dynamics with NAMD. Journal of Computational Chemistry 2005, 26, 1781–1802.

44. Huang, J.; Rauscher, S.; Nawrocki, G.; Ran, T.; Feig, M.; de Groot, B. L.; Grubmüller, H.; MacKerell Jr, A. D. CHARMM36m: an improved force field for folded and intrinsically disordered proteins. Nature methods 2017, 14, 71–73.

45. Klauda, J. B.; Venable, R. M.; Freites, J. A.; O’Connor, J. W.; Tobias, D. J.; Mondragon-Ramirez, C.; Vorobyov, I.; MacKerell, A. D.; Pastor, R. W. Update of the CHARMM All-Atom Additive Force Field for Lipids: Validation on Six Lipid Types. Journal of Physical Chemistry B 2010, 114, 7830–7843.

46. Jorgensen, W. L.; Chandrasekhar, J.; Madura, J. D.; Impey, R. W.; Klein, M. L. Comparison of simple potential functions for simulating liquid water. The Journal of Chemical Physics 1983, 79, 926–935.

47. Jo, S.; Kim, T.; Im, W. Automated builder and database of protein/membrane complexes for molecular dynamics simulations. PLoS ONE 2007, 2.

48. Reid, J. K. Large Sparse Sets of Linear Equations; Academic Press: London, 1971; pp 231–254.

49. Martyna, G. J.; Tobias, D. J.; Klein, M. L. Constant pressure molecular dynamics algorithms. J. Chem. Phys. 1994, 101, 4177–4189.

50. Feller, S. E.; Zhang, Y.; Pastor, R. W.; Brooks, B. R. Constant pressure molecular dynamics simulation: The Langevin piston method. J. Chem. Phys. 1995, 103, 4613– 4621.

51. Humphrey, W.; Dalke, A.; Schulten, K. VMD: Visual molecular dynamics. Journal of Molecular Graphics 1996, 14, 33–38.

52. Immadisetty, K.; Polasa, A.; Shelton, R.; Moradi, M. Elucidating the Molecular Basis of Spontaneous Activation in an Engineered Mechanosensitive Channel. Computational and Structural Biotechnology Journal 2022,

53. Immadisetty, K.; Hettige, J.; Moradi, M. What Can and Cannot Be Learned from Molecular Dynamics Simulations of Bacterial Proton-Coupled Oligopeptide Transporter GkPOT? J. Phys. Chem. B 2017, 121, 3644–3656.

54. Immadisetty, K.; Hettige, J.; Moradi, M. Lipid-Dependent Alternating Access Mechanism of a Bacterial Multidrug ABC Exporter. ACS Central Science 2019, 5, 43–56.

55. Bakan, A.; Meireles, L. M.; Bahar, I. ProDy: Protein dynamics inferred from theory and experiments. Bioinformatics 2011, 27, 1575–1577.

56. Govind Kumar, V.; Ogden, D. S.; Isu, U. H.; Polasa, A.; Losey, J.; Moradi, M. Prefusion Spike Protein Conformational Changes Are Slower in SARS-CoV-2 than in SARS-CoV-1. Journal of Biological Chemistry 2022, 298.

57. Immadisetty, K.; Polasa, A.; Shelton, R.; Moradi, M. Elucidating the Molecular Basis of pH Activation of an Engineered Mechanosensitive Channel. bioRxiv 2021,

58. Polasa, A.; Mosleh, I.; Losey, J.; Abbaspourrad, A.; Beitle, R.; Moradi, M. Developing a Rational Approach to Designing Recombinant Proteins for Peptide-Directed Nanoparticle Synthesis. Nanoscale Adv. 2022,

59. Moradi, M.; Enkavi, G.; Tajkhorshid, E. Atomic-level characterization of transport cycle thermodynamics in the glycerol-3-phosphate:phosphate transporter. Nat. Commun. 2015, 6, 8393.

60. Moradi, M.; Sagui, C.; Roland, C. Calculating relative transition rates with driven nonequilibrium simulations. Chem. Phys. Lett. 2011, 518, 109–113.

61. Moradi, M.; Sagui, C.; Roland, C. Invstigating rare events with nonequilibrium work measurements: I. nonequilibrium transition path probabilities. J. Chem. Phys. 2014, 140, 034114.

62. Moradi, M.; Sagui, C.; Roland, C. Invstigating rare events with nonequilibrium work measurements: II. transition and reaction rates. J. Chem. Phys. 2014, 140, 034115.

63. Moradi, M.; Tajkhorshid, E. Mechanistic Picture for Conformational Transition of a Membrane Transporter at Atomic Resolution. Proc. Natl. Acad. Sci. USA 2013, 110, 18916–18921.

64. Ogden, D.; Moradi, M. Structure and Function of Membrane Proteins; Springer US: New York, NY, 2021; pp 289–309.

65. Fiorin, G.; Klein, M. L.; Hénin, J. Using collective variables to drive molecular dynamics simulations. Mol. Phys. 2013, 111, 3345–3362.

66. Chen, M.; Samuelson, J. C.; Jiang, F.; Muller, M.; Kuhn, A.; Dalbey, R. E. Direct interaction of YidC with the Sec-independent Pf3 coat protein during its membrane protein insertion. Journal of Biological Chemistry 2002, 277, 7670–7675.

67. Yu, Z.; Koningstein, G.; Pop, A.; Luirink, J. The conserved third transmembrane segment of YidC contacts nascent Escherichia coli inner membrane proteins. Journal of Biological Chemistry 2008, 283, 34635–34642.

68. Harkey, T.; Govind Kumar, V.; Hettige, J.; Tabari, S. H.; Immadisetty, K.; Moradi, M. The Role of a Crystallographically Unresolved Cytoplasmic Loop in Stabilizing the Bacterial Membrane Insertase YidC2. Scientific Reports 2019, 9, 14451.

69. Kumazaki, K.; Tsukazaki, T.; Nishizawa, T.; Tanaka, Y.; Kato, H. E.; Nakada-Nakura, Y.; Hirata, K.; Mori, Y.; Suga, H.; Dohmae, N. et al. Crystallization and preliminary X-ray diffraction analysis of YidC, a membrane-protein chaperone and insertase from Bacillus halodurans. Acta Crystallographica Section F:Structural Biology Communications 2014, 70, 1056–1060.

